# Maternal age and genome-wide failure of meiotic recombination are associated with triploid conceptions in humans

**DOI:** 10.1101/2025.02.15.637872

**Authors:** Ludovica Picchetta, Christian Simon Ottolini, Xin Tao, Yiping Zhan, Vaidehi Jobanputra, Carlos Marin Vallejo, Francesca Mulas, Elvezia Maria Paraboschi, Maria José Escribá Pérez, Thomas Molinaro, Christine Whitehead, Pavan Gill, Emily Mounts, Dhruti Babariya, Laura Francesca Rienzi, Filippo Maria Ubaldi, Juan Antonio Garcia-Velasco, Antonio Pellicer, Shai Carmi, Eva R Hoffmann, Antonio Capalbo

## Abstract

Triploid and haploid conceptions are not viable and are a common occurrence in humans, where they account for 10% of all pregnancy losses. Despite the parent-of-origin being important in the etiology of the pregnancy, our knowledge of their causes is limited, especially at the point of conception. Using a dataset of 96,660 biopsies and a validation dataset of 44,324 from human blastocysts embryos generated by intra-cytoplasmic sperm injection (ICSI), we estimate that 1.1% of human conceptions (n=1,063) contain extra or missing chromosome sets in zygotes. We identify a maternal age effect, with a 1.059 per year increased risk in triploidy/haploidy (p=0.0008). In 0.03% of couples, we identified three or more triploid/haploid embryos, suggesting a personal risk effect (p=0.03). Genotype analysis of 55 triploid embryo biopsies and their parents show that one third of maternal triploid conceptions originate in meiosis I and two thirds in meiosis II. Seven of these embryos are inferred to have entirely failed to initiate meiotic recombination genome-wide, suggesting that human oocytes with pervasive meiotic recombination failure that are formed during fetal development are capable of ovulation in adult life. Finally, we identify a new type of genome-wide maternal isodiploidy (two maternal chromosome sets) in 0.05% of embryos (41 of 74,009). Collectively, our findings shed light on the biology of meiosis and the formation of human oocytes with the number of chromosome sets.

## Introduction

Meiosis is a specialized cell division that reduces the genomic content by one half, such that when oocyte and sperm combine at fertilization, the diploid content is restored in the offspring. Failure to reduce the diploid content results in additional (polyploidy) or missing (haploidy) chromosome sets, a life-limiting condition in humans and other animals (1,2).

In humans, 1% of conceptions are estimated to be triploid and they make up 10% of all pregnancy losses, including during IVF treatment (3–8). As early as 1967, ploidy abnormalities were observed in tissue from pregnancy losses, chorionic carcinoma, and molar pregnancies, and they are recognized as one of the leading causes of pregnancy complications (9,10). Genetic analysis of fetal losses shows that about one third of maternal-origin (digynic) triploidy originate in meiosis I (with the remaining in meiosis II), and that paternal (diandric) triploidy is mainly due to fertilization by two sperm or diploid single sperm (11). The parent-of-origin of the error is important clinically as it is associated with distinct maternal risks, fetal abnormalities, and prenatal sonographic features (12–14).

Triploidy was previously thought to be a random occurrence of abnormal fertilization (3,15–18). However, recent data from mouse models (19) and genetic analysis of human polyploid conceptions suggest that a subset of individuals may be predisposed to abnormal ploidy conceptions (20–23). Epidemiological studies suggest no association with parental age or aberrant meiotic recombination (24,25), two factors strongly associated with aneuploidy in oocytes (where only one or a few chromosomes are affected). Triploidy is also common in other animals such as horses (1) and salmon (2).

Our current knowledge of polyploid conceptions is limited to studies at stages of post-implantation fetal development, where selection against dysfunctional genomic constellations has already occurred. Therefore, we set out to leverage information from large datasets of preimplantation embryos that underwent genetic testing following in vitro fertilization (IVF) and extended culture to the blastocyst stage of embryo development. This allowed us to detect a maternal age effect, genome-wide recombination failure, as well as a new type of isodiploidy that affects human oocytes.

## RESULTS

### Incidence of ploidy abnormalities in ICSI-generated blastocysts

To determine the incidence of ploidy abnormalities in human preimplantation embryos, we used data obtained from 96,660 trophectoderm biopsies (DATASET A; see Methods and Supplementary Table 1) from 20,187 IVF cycles that used a targeted next-generation sequencing (NGS) protocol for preimplantation genetic testing for aneuploidy (PGT-A) or monogenic disorders (PGT-M) (Figure 1). All embryos generated used intracytoplasmic sperm injection (ICSI) and two pronuclei were identified after fertilization (2PN) during the routine, static, morphological assessment performed by the embryologists. 93,986 embryos provided high quality informative samples of which 66,993 embryos were euploid (71.3%; 95% CI: 71.0-71.6%) and 25, 930 were aneuploid (27.6%; 95% CI: 27.3-27.9%) (Figure 1 and Supplementary Figure 1A). As expected, the aneuploidy rate was positively correlated with advanced maternal age, regardless of the indication for testing (PGT-A or PGT-M) (Supplementary Figure 2).

**Figure 1.**
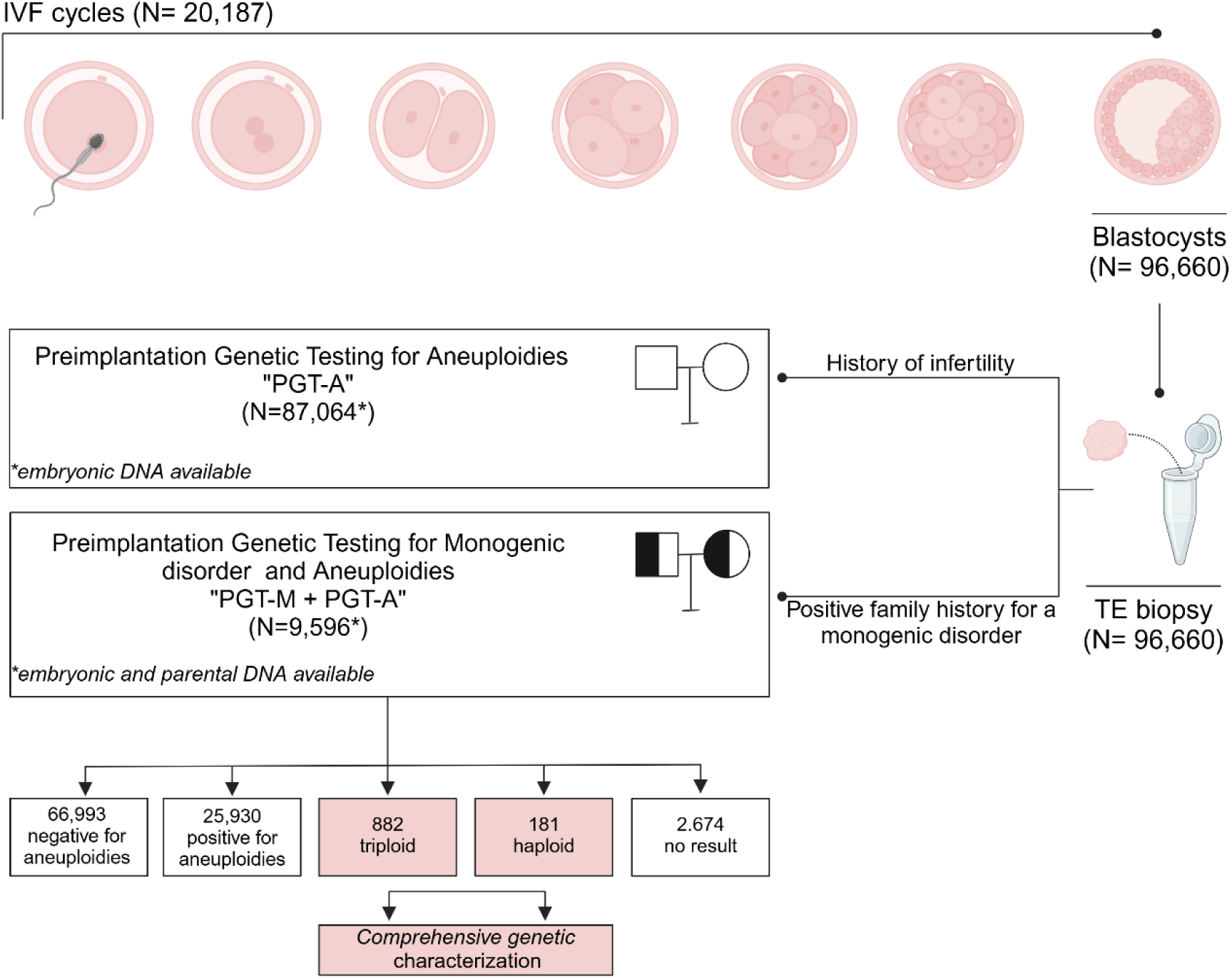
Summary of the main dataset and clinical workflow. A total of 96,660 blastocyst stage embryos, derived from 20,187 IVF cycles with apparently normal fertilization, underwent trophectoderm biopsy for preimplantation genetic testing. Among these, 87,064 TE biopsies were of embryos from couples with infertility and PGT was intended to detect aneuploidy (PGT-A). Conversely, in 9,596 TE biopsies from couples with a positive family history for a monogenic disorder, PGT was performed for both aneuploidy and the specific monogenic disorder (PGT-M). In this subgroup, both embryonic and parental DNA were available for genetic analysis. Comprehensive genetic characterization was performed on triploid and haploid samples.

We discovered ploidy abnormalities in 1.1% of embryos (n=1,063/93,986; 95% CI: 1.06-1.20%). Of these, 181 (17.0%; 95% CI: 14.8-19.4%) were found to be haploid (Figure 1 and Supplementary Figure 1A). Only one haploid embryo had a Y chromosome, while the X chromosome was present in all the others, demonstrating that haploidy is almost entirely representative of paternal origin (p<2.2×10^-16^; binomial exact test; Supplementary Figure 1B).

Among the 1,063 embryos with ploidy abnormalities, 882 were triploid (83.0%; 95 CI: 80.6-85.2%; Figure 1 and Supplementary Figure 1A), indicating a prevalence nearly five times greater compared to haploid abnormalities in blastocysts. Among the triploid embryos, 433 had three X chromosomes (69, XXX; 49.1%); 433 embryos contained two X chromosomes and one Y (69, XXY; 49.1%), and 16 embryos had one X chromosome and two Y (69, XYY; 1.8%) (Supplementary Figure 1B).

These observations suggest a 4.8-fold preponderance of triploid over haploid blastocysts (882 compared to 181). Thus, the excess in prenatal diagnosis of triploid fetal losses compared to haploid is not only due to selection during fetal development, but a bias at the blastocyst stage.

### Parental and meiotic phase of origin of ploidy abnormalities

Next, we investigated the parental contribution and meiotic origin of ploidy errors in the 1,063 embryos that contained ploidy abnormalities. We first used a modeling approach, based on the sex chromosome constellation (XXX/XXY/XYY), and assumed independence between parental and meiotic origins. We estimated that in 94.6% of triploid embryos, the extra chromosome set was of maternal origin, whereas 5.4% were of paternal origin (Supplementary Figure 1C). Triploid embryos arose from failure to segregate chromosomes both in the first and second meiotic division (33.3% and 66.7%, respectively; Supplementary Figure 1D). In contrast, 98.9% of the 181 haploid embryos were due to a missing set of paternal chromosomes, based on having only a single haploid embryo with a Y chromosome (Supplementary Figure 1C).

To directly determine the parent-of-origin and meiotic phase at which entire chromosome sets failed to segregate, we used single nucleotide polymorphism (SNP) genotyping data from embryos and parents (‘trios’) undergoing testing for monogenic disorders. Of 55 trios analysed, haploid embryos were confirmed to be predominantly caused by an absence of paternal genome (12 of 14; 85.7%; 95% CI: 57.2-98.2%). All 41 triploid embryos (100%; 95% CI: 91.4-100.0%) were due to female meiosis errors.

Meiotic recombination is reduced near centromeres and informative SNPs (within 5cM from the centromere) can be used to detect the presence of homologs (meiosis I) versus sister chromatids (meiosis II) in triploid embryos. Eleven of the 41 triploid embryos (26.8%; 95% CI: 14.2-42.9%) had both maternal homologs for all chromosomes and were therefore caused by genome-wide segregation failure in meiosis I. The remaining embryos carried alleles from only a single homolog around all centromeres and were inferred to originate from genome-wide failure to segregate sister chromatids at meiosis II (n=30/41; 73.2%; 95% CI: 57.1-85.8%). Triploid conceptions that originate from maternal meiosis are also found in natural conceptions (11,26,27), suggesting that these observed effects are not due to ovarian stimulation.

### Recombination rates of triploid embryos

To investigate a potential causal relationship between ploidy abnormalities and recombination events, we analyzed SNP inheritance patterns for the 41 triploid embryos for which parental genotyping data was available. Meiotic recombination between homologous chromosomes caused switches between regions of single parental homolog (SPH) or biparental homolog (BPH) along chromosomes. We detected the switches using the delta inconsistency score (Methods) after correcting for the number of usable SNPs (Supplementary Table 2, Supplementary Figure 3).

The number of detectable recombination events ranged from 0 to 26 in the triploid embryos of maternal origin (Figure 2 A-B, Supplementary Table 2). Of these, most contained between 8 to 26 recombination events genome-wide (Figure 2C). This number of recombination events is lower than the 41-46 previously observed in human preimplantation embryos (28,29), particularly that the number of SPH/BPH switches is higher than the number of recombination events that would have been observed in a diploid embryo (Supplementary Note 1). This discrepancy may serve as a potential biomarker for gross chromosomal abnormalities. However, a possible bias may result from differences between the two technologies as some crossovers may be missed due to the sparse genome coverage of our assay (e.g., Figure 2C).

**Figure 2.**
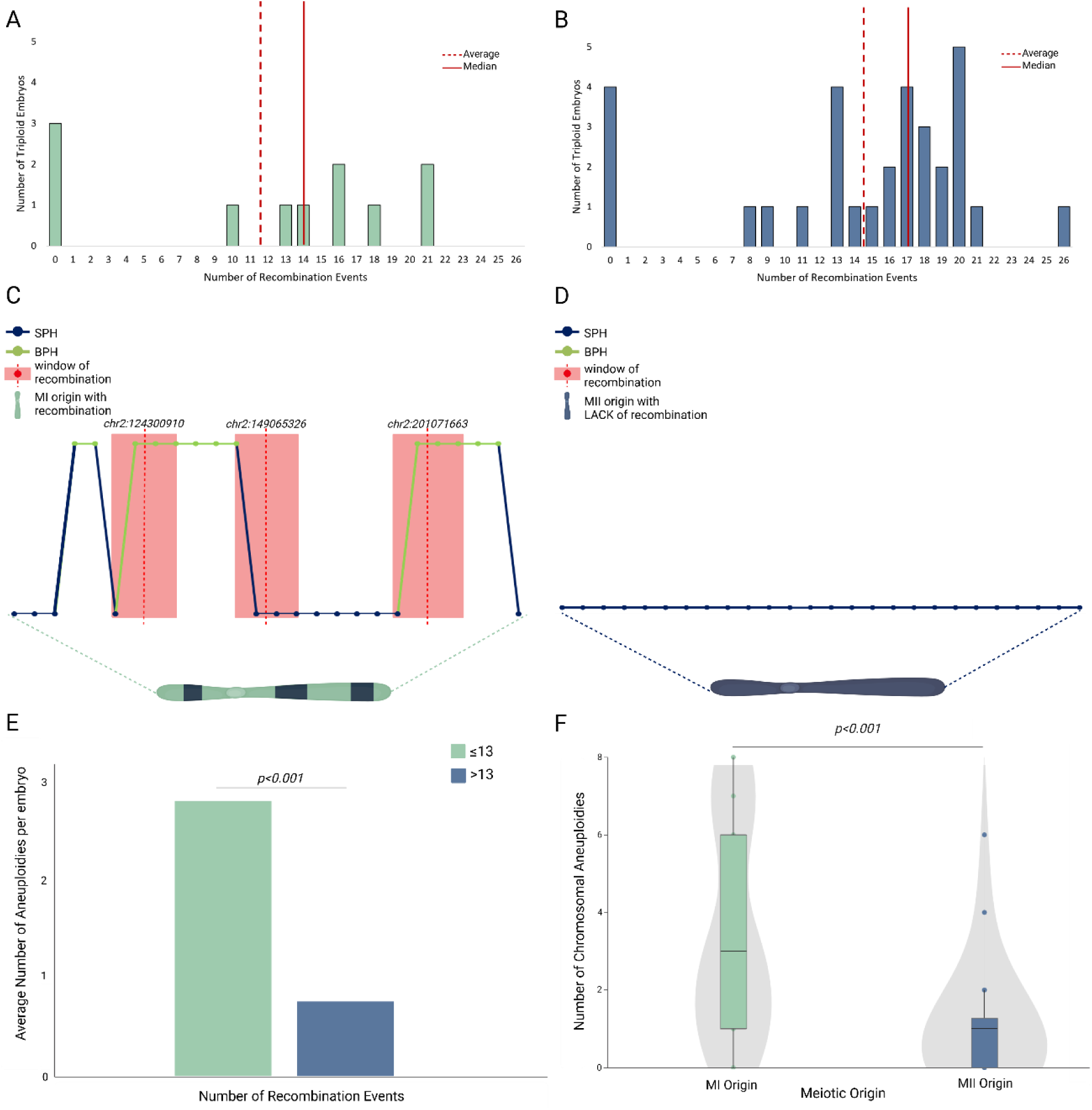
Recombination events in triploid embryo and their association with aneuploidy rate. (A-B) Bar plots depicting recombination events in triploid embryos of meiosis I origin (A) and meiosis II origin (B). (C-D). Graphic representation of recombination windows detected in chromosome 2 from an embryo with an average number of recombination events and meiosis I origin (C) and an embryo with lack of genome wide recombination and meiosis II origin (D). Each dot represents the delta score for each SNP, with green indicating a BPH state for maternal alleles from a meiosis I error, and blue indicating an SPH state for maternal alleles from a meiosis II error. The red dot corresponds to a point of switch of the average score. The red rectangle highlights the window of recombination described as the area within the identified point of switch and the closest preceding SNPs with an opposite average score value. (E) Bar plot illustrating a statistically significant correlation between the average number of aneuploidies per embryo and recombination rates, comparing one group of embryos with low recombination rates (<13 events) and a second group with higher recombination rate (>13 events); 13 is the mean number of recombination events in our dataset. (F) Box plot and violin plots showing a different rate of additional aneuploidies in triploid embryos of meiosis I origin (green) and meiosis II origin (blue).

Seven of the triploid embryos were inferred to lack crossovers genome-wide (7/41 or 17.1%; 95% CI: 7.1-32.1%; Figure 2D). A permutation test was employed to test if the lack of genome-wide recombination could be interpreted as an “expected” finding and revealed that these embryos constituted an outlier group (P<0.001) compared to the remaining embryos. Failure to initiate meiotic recombination was inferred in diploid oocytes originating both from meiosis I and meiosis II (Figure 2A and B). Genome-wide recombination failure has previously been inferred in one human oocyte (28) and shown to be associated with single chromosome aneuploidies. Since all fertilized oocytes contained a polar body for the ICSI procedure, our observations suggest that failure to segregate one chromosome set into the polar body is another consequence of genome-wide recombination failure.

We detected additional chromosomal aneuploidies in triploid embryos. This finding was statistically correlated with meiotic origin of triploidy, with triploid embryos of MI origin being more prone to additional segregation errors (P<0.001) (Figure 2F). Furthermore, a lower number of genome-wide recombination events was also found to be statistically correlated with a higher number of chromosomal aneuploidies in the triploid embryos (P<0.001) (Figure 2E).

### Recombinant genome-wide uniparental isodisomy

The lack of paternal genomes in the haploid embryos and the prevalence of two maternal chromosome sets in the triploid embryos predict a class of embryos that are diploid but contain only maternal chromosome sets (maternal isodiploidy).

To detect this, we developed an algorithm (Methods) that mapped recombinant sister chromatids, genome-wide, after segregation failure in meiosis II and a normal meiosis I. We detected 60 such events amongst 74,009 embryos from DATASET D (see Methods and Supplementary Table 1). Manual checking of PGT plots confirmed the fidning for 41 of these (0.05%,95% CI:0.04-0.07%) (Figure 3A and B). We refer to these as ‘recombinant isodiploidy’ and they were characterized by homozygosity in pericentromeric regions, but with few detectable heterozygosity regions throughout the rest of the genome, indicating that crossovers occurred between the two parental homologs during prophase I. Additionally, fingerprinting analysis confirm a loss of genetic correlation between the affected embryo and the paternal DNA sample (Figure 3C) as well as with corresponding sibling embryos. The latter is to be expected due the absence of one parental genome in the affected embryo compared to the rest of sibling embryos sharing genetic similarities in both the maternally and paternally inherited genomes.

**Figure 3.**
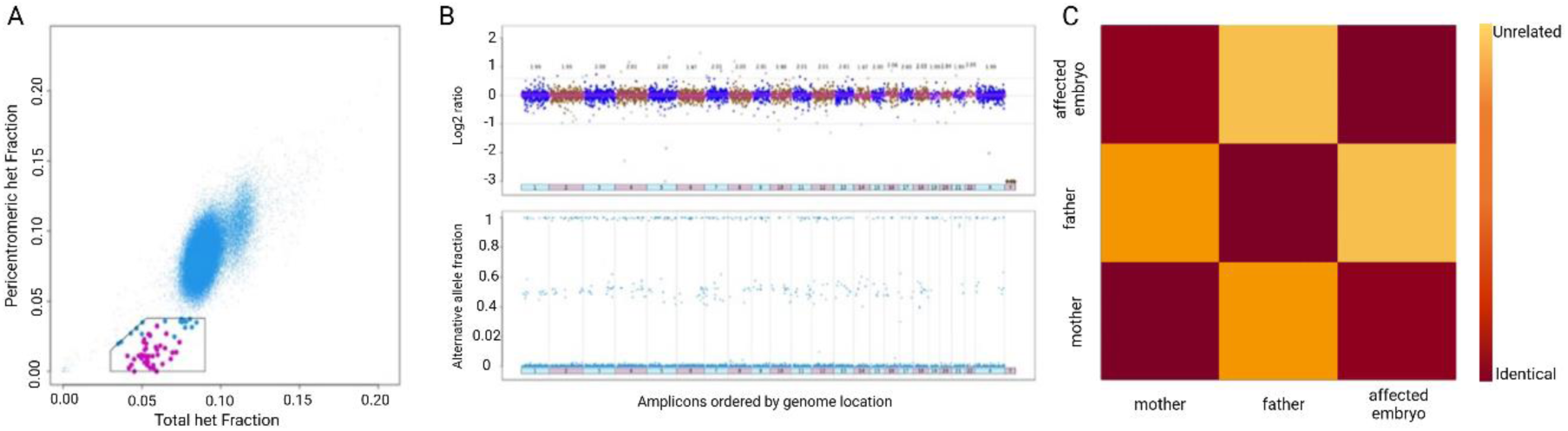
Recombinant genome-wide uniparental isodisomy in preimplantation embryos. (A) Distribution of 74,009 samples according to their total het fraction (Y axis) and pericentromeric het fraction (X axis). Dots in the pentagonal box represent putative isodiploid embryos, magenta-colored dots are embryos confirmed to be isodiploid by PGT and fingerprinting plots. (B) PGT plots of an affected embryo. Top: the amplicons distribution in terms of copy number. Bottom: the alternative allele fraction, showing regions of loss of heterozygosity (absent dots with values between 0.1 and 0.9). (C) Fingerprinting plot of an affected embryo and its parents. Dark red corresponds to genetic identity, while the lighter the color the lower the genetic correlation.

### Parental Age correlation

The large size of our dataset, which include 1,063 ploidy abnormalities, allowed us to correlate their incidence with parental age. The mean maternal age was 35.65 years (± 2.9 years; range 20-47) and the mean paternal age was 37.22 (± 3.03 years; range 19-79). We detected a positive and statistically significant correlation (OR=1.046 per year; p<0.001) for ploidy abnormalities and maternal age (Figure 4). Triploidy had the strongest correlation with maternal age (OR=1.059 per year; p<0.001) (Figure 4), consistent with the finding that they have a predominant maternal origin in our dataset. The risk of having a triploid conception at female age 40 was 76% higher than at age 30.

**Figure 4.**
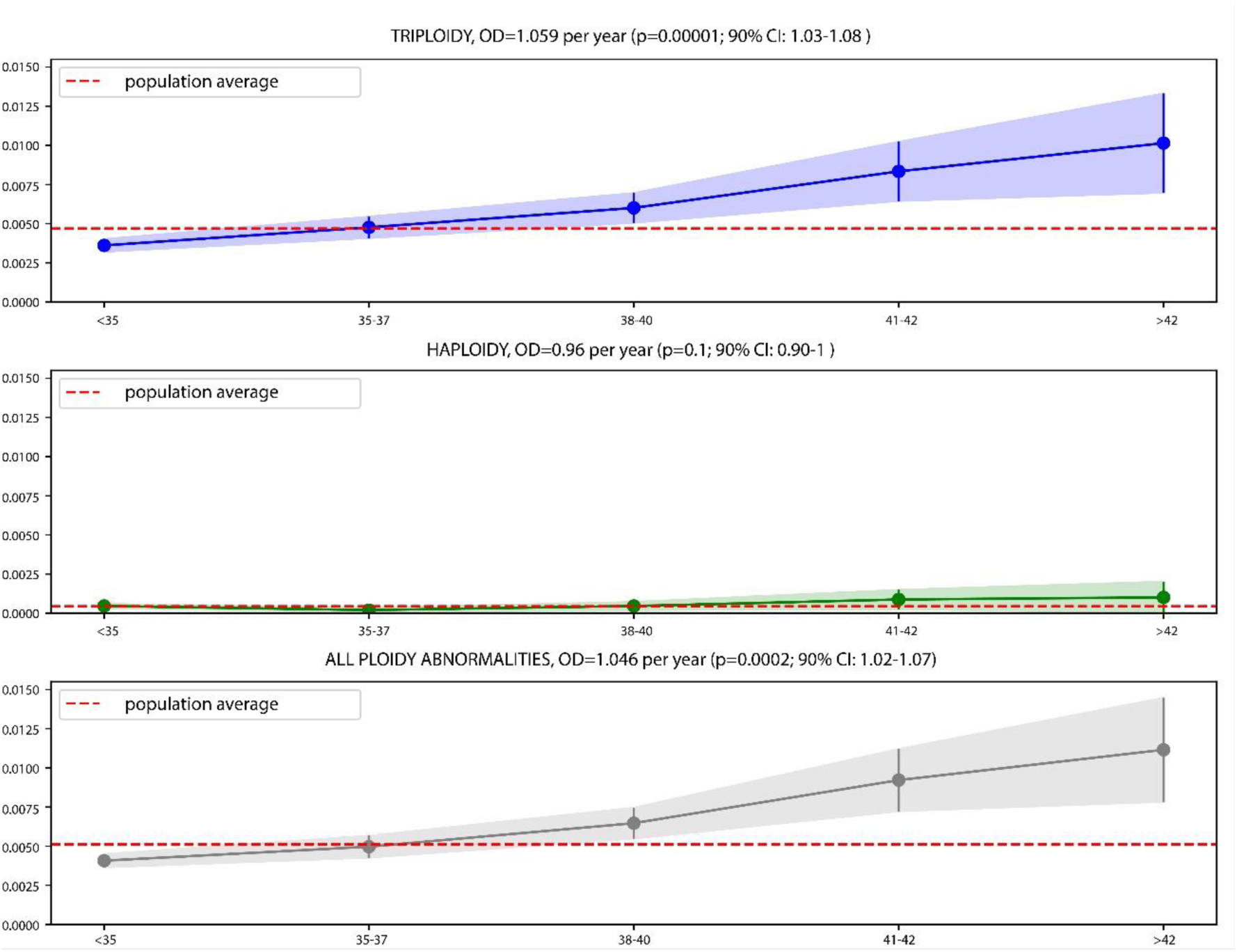
The risk of ploidy abnormalities rises with advancing maternal age. Plots illustrating the relationship between maternal age and various subsets of ploidy abnormalities. Maternal ages were classified using the SART age groups (X axis). The Y axis shows the fraction of affected embryos: triploid, haploid, and all ploidy abnormalities.

Paternal age was significantly associated with an increased risk of triploid conceptions in the univariate analysis. However, this association disappeared in the multivariate analysis, as paternal age was linked to maternal age (Supplementary Table 3).

The association between maternal age and triploidy was confirmed in an independent dataset of 44,324 embryos (DATASET B; see Methods and Supplementary Table 1), where we observed an OR= 1.03 per year (p=0.03, Supplementary Figure 4). We conclude that maternal age is associated with diploid oocyte formation and triploid blastocysts.

### Maternal age is associated with elevated risk of abnormal fertilization

We noted a slight difference in the effect size of female age in the two datasets, as well as a slightly lower frequency of triploidy in DATASET B (0.7%; 293 of 44,324; 95%CI: 0.6-0.74). We considered two, not mutually exclusive, explanations, including that the smaller sample size of the second data set might explain the differences. However, a second explanation may be that the difference is caused by fewer fertilized oocytes being misclassified in the second dataset as being 2PN, when in fact they were 3PN, since time lapse incubators were used. Zygotes with 3PN are known to have a higher incidence of triploidy and may have contributed to the higher incidence in the larger dataset, where the latter did not use time lapse incubators (30).This predicts that abnormal fertilization should be elevated with maternal age.

To test whether abnormal fertilization events were increased with maternal age, we used DATASET C (see Methods and Supplementary Table 1) comprising 93,341 zygotes for which pronuclei number was available. Advanced maternal age was statistically significantly associated with an increased risk of abnormal fertilization following ICSI (1PN and >2PN; p=1.21×10^7^; Negative binomial regression). In particular, the presence of three pronuclei had the strongest correlation with maternal age (p=2×10^-8^; Supplementary Figure 5). Thus, the higher chance of exclusion of 3PN fertilized oocytes in the smaller validation dataset, misses a subset of maternal triploid conceptions that would have arisen from abnormally pronucleated zygotes.

### Recurrence of ploidy abnormalities in a subset of patients

To determine whether some patients are at an increased risk of recurrent ploidy abnormalities, we analysed the number of affected embryos in any given treatment cycle. We used the original DATASET A consisting of 96,660 embryos obtained from 20,187 cycles and 20,187 different couples. We found that 19,288 cycles produced no embryos with ploidy level defects, 852 cycles produced one embryo with a ploidy level defect (4.2%; 95% CI: 3.9-4.5%), 41 cycles produced two (0.2%; 95% CI: 0.1-0.3%), and six cycles produced three (0.03%; 95% CI: 0.01-0.06%) (Supplementary Table 4). In four of the six couples with three ploidy abnormal embryos, the maternal age was <35 years. Under a null hypothesis of no family-specific causes of ploidy abnormalities, the only factor affecting the ploidy occurrence is maternal age and embryos with ploidy abnormalities should therefore be randomly distributed among embryonic cohorts from women of similar maternal ages. However, this was not the case, as a permutation analysis showed that the recurrence of three or more ploidy abnormal embryos from the same couple cannot be explained as a random event (Figure 5). Therefore, this analysis suggests there may be other factors increasing the risk of ploidy abnormalities recurrence.

**Figure 5.**
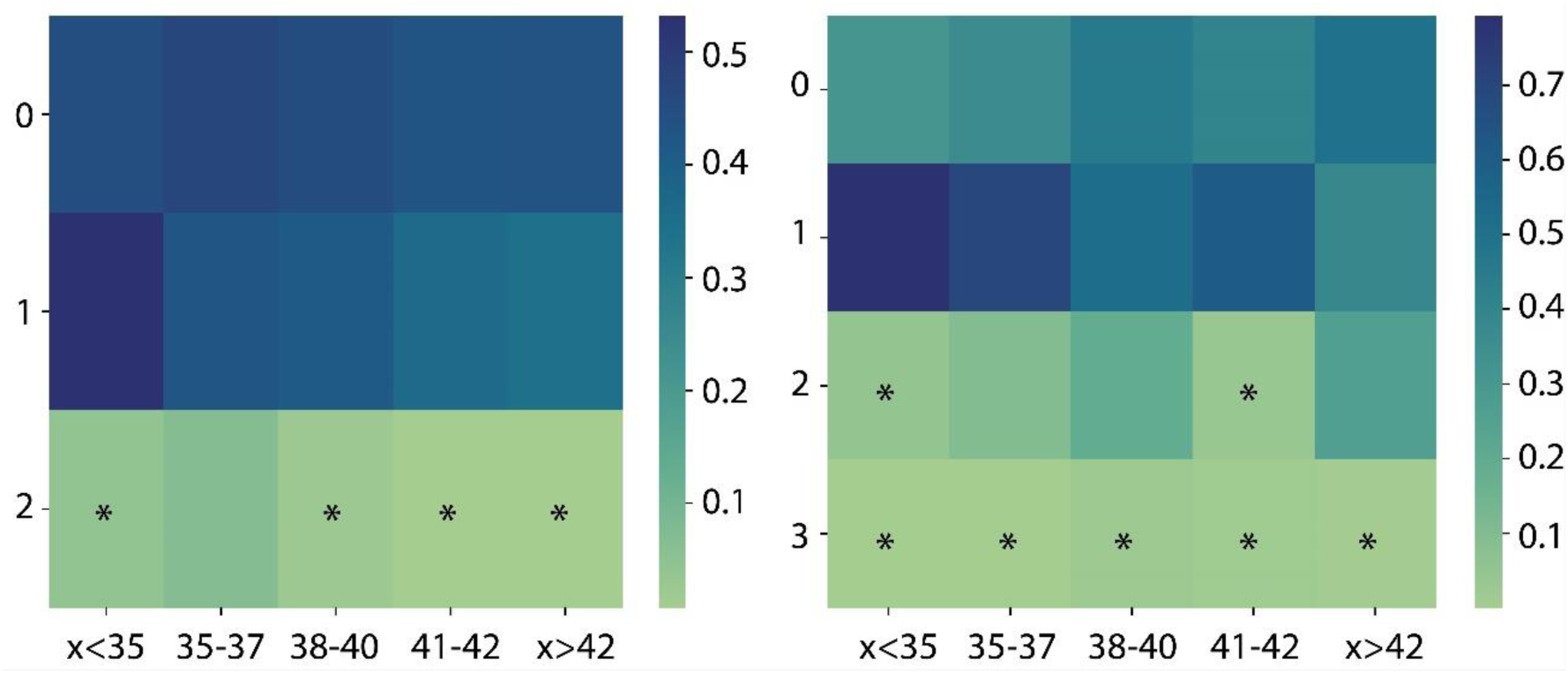
Unexpected per-family ploidy error recurrence. Heatmap of Bonferroni adjusted P-values obtained for each maternal age category (columns) and each ploidy error recurrence count (rows) for haploidy (left) and triploidy (right). P-values represent the probability of observing the event by random chance. Heatmap’s squares marked with “*” were considered statistically significant (p<0.05; Bonferroni adjusted).

## DISCUSSION

In this study, we investigated the parental and meiotic origins of haploid and triploid blastocysts in humans. We used several independent large datasets from PGT cycles where ploidy abnormalities can be detected. The use of preimplantation human embryos created *in vitro* following ICSI in infertility patients using PGT-A and non-infertility patients using PGT-M allowed us to indirectly infer if a gamete was affected by a ploidy abnormality. The overall incidence of ploidy abnormalities inferred as originating from gametes was 1.15%, with triploidy accounting for the vast majority (83%) (31). These rates are similar from those reported in prenatal diagnosis of natural conceptions (26), when the lack of dispermy in our dataset is considered.

Some of our findings were unexpected, including that the haploid conceptions were mainly of paternal origin, indicative of sperm without a genome or lack of genome decondensation at fertilization. Consistent with this, oocytes were more likely to be diploid than nulliploid, suggesting that when genome-wide chromosome segregation fails, the chromosomes are more likely to remain in the oocyte rather than segregate to the polar body. The robust datasets also allowed us to detect a maternal age effect on generation of diploid oocytes at both the levels of meiosis I and meiosis II, as well as on abnormal fertilization. These effects are unlikely to be due to ovarian stimulation and ICSI, respectively, since our data show important similarities to those of in vivo conceptions, including the higher incidence of errors leading triploidy in meiosis II compared to meiosis I (26,32).

It is long known that meiotic recombination may fail on individual chromosome pairs, especially the small chromosomes 21 and 22 (33), resulting in an elevated risk of aneuploid and trisomic conception (e.g., trisomy 21). Our findings, however, reveal genome-wide recombination failure that results in diploid oocytes at meiosis I and meiosis II. Meiotic recombination is initiated by the highly conserved Spo11-Top6BL transesterase during meiotic prophase I (34–40). Recent findings in budding yeast, however, reported that deletion of *spo11* in some cases resulted in diploid spores, without evidence of recombination (41). Deletion of *spo11* causes both male and female infertility in mouse (42,43), due to checkpoints that eliminate germ cells that fail to engage in meiotic recombination, synapsis and silencing. The meiotic silencing checkpoint, however, is sexually dimorphic with a less stringent response in females (44). Furthermore, BLC-2 has been shown to eliminate recombination-defective oocytes in mouse (45). Our findings suggest that initiation of meiotic recombination can fail genome-wide in human fetal oocytes, and that at least some of these may escape checkpoints during the fetal stages and are capable of ovulating decades later.

Failure to segregate sister chromatids during meiosis II may be the main cause of triploid conceptions in humans. Meiosis II failure also resulted in recombinant isodiploid blastocytes caused by diploidy in the oocyte and lack of sperm DNA. These errors may lead to extreme imprinting disorders affecting more than 150 genes (46). Fertilization abnormalities were also affected by advancing maternal age, indicating oocytes become more error-prone with age in terms of segregating sister chromatids as well as forming 3PN. The causes of such effects remain to be investigated; however, our findings suggest that genome-wide failure in meiotic recombination during fetal life, together with meiosis I and meiosis II segregation failure in adult life, result in ploidy abnormalities in human oocytes.

## METHODS

### Study design

A retrospective, cohort study was conducted by Juno Genetics, a clinical laboratory improvement amendment (CLIA)–certified genetic testing laboratory assessing embryos from 62 referring reproductive medicine clinics in the United States between January 2020 and September 2023. Raw genetic data, collected as standard clinical practice, were leveraged for nonclinical analysis of ploidy level abnormalities and any potential association with clinical/embryological parameters. To evaluate the impact of parental age, patients were classified in five distinct age groups based on SART classification (49).

The first part of the study aimed at the comprehensive characterization of haploidy and triploidy. This analysis was conducted on a cohort drawn from a total of 96,660 trophectoderm biopsy samples taken from blastocyst stage embryos derived from zygotes displaying two pronuclei following in vitro fertilization (IVF) via intracytoplasmic sperm injection (ICSI) (20,187 cycles) (referred to as DATASET A; Supplementary Table 1). All embryos were subjected to clinical preimplantation genetic testing (PGT) for either chromosomal aneuploidy alone (PGT-A) due to history of infertility (n=87,064) or concurrent aneuploidy testing and testing from monogenic disorders (PGT-M) (n= 9596) due to a positive family history for a monogenic disorder. PGT was performed using a targeted next generation sequencing platform (47,48). To validate the findings on ploidy abnormalities and their relationship with clinical factors (e.g., maternal age) an independent cohort of 44,324 trophectoderm biopsy samples taken from blastocyst stage embryos (referred to as DATASET B; Supplementary Table 1) were analyzed. Additionally, the association between maternal age and ploidy abnormality was tested using DATASET C (Supplementary Table 1), a cohort comprising 93,341 zygotes with corresponding annotation of the number of pronuclei displayed around 16 hours post in vitro fertilization. Finally, a fourth database (referred to as DATASET D; Supplementary Table 1) of 74,009 embryo biopsies with a 46,XX karyotype tested for aneuploidy by targeted NGS between January 2020 and September 2023, and partially overlapping with DATASET A, was used to investigate the incidence and origin of isodiploidy.

IRB approvals were obtained prior to research data analysis (Advarra Pro00074493; WIRB 1053149 and CEIM - HOSPITAL UNIVERSITARIO Y POLITÉCNICO LA FE N° 2024-0405-1).

### Sequencing data analysis

PGT was performed clinically for embryo biopsy samples by first performing a targeted DNA amplification step followed by next generation sequencing (NGS) by PGTseq (PGTseq Technology Inc, Basking Ridge, New Jersey) analysis (47,48). In short, NextSeq500/550 or NovaSe6000 Mid and High Output Kit v2.5 NGS-based PGT-A was used for TE biopsy chromosome copy number analysis, with around 5,000 amplicons and SNPs per sample. Proprietary PGTseq software was used for bioinformatics and automatic calls of chromosome copy number as previously described (25, 30). Embryo ploidy abnormalities were clinically reported following a validated analytical pipeline of combined sequencing quantitation and genotyping allele ratio data analysis. Briefly, the allele ratio was obtained for each heterozygous SNP in each biopsy sample. In the instance of a balanced chromosome copy number and unbalanced allele ratios (2:1) across the genome, indicative of 3 copies for each chromosome, the embryo was deemed triploid. If there was a total loss of heterozygosity across the genome, indicating the presence of only one allele with an allele ratio of 1:0 per chromosome, the embryo was deemed haploid.

A bespoke research bioinformatic pipeline was developed to more comprehensively analyze raw sequencing data from biopsy samples and to further investigate preimplantation embryo ploidy anomalies. BAM files were aligned against the GRCh37 human reference using BWA (Burrows Wheeler Aligner) and FREEBAYES v1.3.2 was used with default settings to identify variants. Using vcf2tsv, genomic VCF files were converted to a tab-separated format suitable for downstream analysis. Customized allele frequency thresholds were established to generate discrete genotype calls based on the proportion of read counts mapped at the specific genomic locus, as follows. Loci with an alternative allele frequency below 0.05 were defined as homozygous reference, loci with allele frequency between 0.2 and 0.8 were deemed heterozygous, whereas loci with allele frequencies higher than 0.95 were called as homozygous reference.

Variants were filtered according to the following criteria: (i) variant was a biallelic single nucleotide polymorphism (SNP) according to gnomAD v2.1.1 (51), allowing higher accuracy of the model; (ii) the depth of coverage was greater than 20x (the average coverage was 360x, ranging from 0 to 10,104x), thus providing sufficient reads to estimate B-allele ratios; (iii) the B-allele frequency (BAF) ranged between 5% and 95%; (iv) variant mapped on an autosome.

### Parental and meiotic phase of origin of ploidy abnormalities using sex chromosome’s copy number values

CN values of the sex chromosomes were used to infer parental origin and meiotic origin of the ploidy abnormality in 1,063 haploid and triploid embryos.

This was accomplished on triploid embryos using a mathematical model based on a priory clinical diagnosis of a ploidy abnormality and the assumption that the error distribution between the two meiotic divisions (i.e., MI, MII) is the same in both sexes:

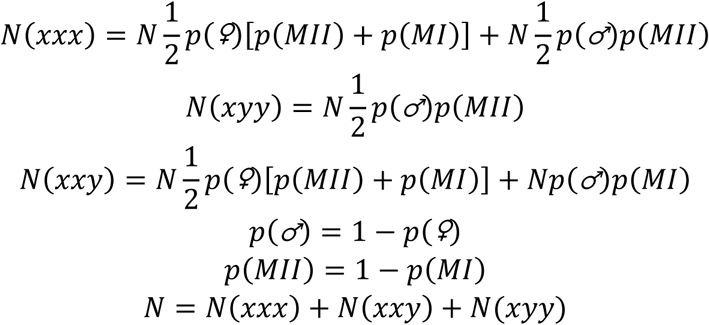

When considering triploid embryos with a ChrX : ChrY copy ratio of 3:0 [N(xxx)], 2:1 [N(xxy)], and 1:2 [N(xyy)], p(♀) is the probability that a triploid is due to a female error, p(♂) is the probability that a triploid is due to a male error, p(MI) is the probability that a triploid is due to a meiosis I error, p(MII) is the probability that a triploid is due to a meiosis II error, and N is the total number of triploid embryos. Therefore, we obtained the following “method-of-moments” estimators:

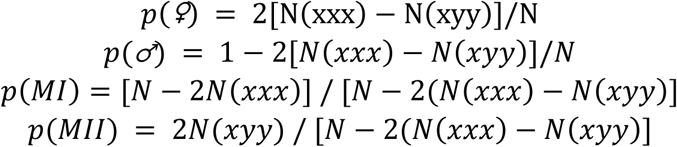

The same approach was also used on haploid samples, following the equation below. Due to the lack of genetic data from the parent in whom the error has occurred, it was not possible to distinguish between meiosis I and meiosis II in this subset.

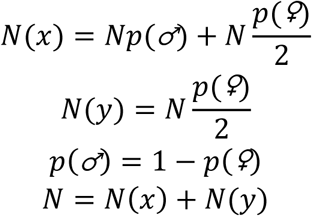

Therefore, we obtain:

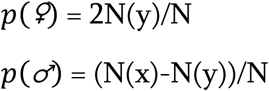

### Parental and meiotic phase of origin of ploidy abnormalities using genotyping data

Using the genotypic output of a subset of ploidy-abnormal embryos analyzed through concurrent PGT-A and PGT-M, a second independent and more comprehensive methodology (compared to sex chromosomes CN analysis) was applied to calculate parent and meiotic origin of the extra/missing haploid set of chromosomes in ploidy-abnormal embryos. In total, embryonic and parental genotyping data from 55 trios were analyzed, including 41 triploids and 14 haploids. To determine the parental origin (in both triploid and haploid abnormal embryos), the informative SNPs had to be opposite homozygous in the two parents. As an example, if the maternal genotype was “AA” and the paternal genotype was “BB” at the same locus, an occurrence of “ABB” for the embryo was defined as a paternal error, whereas “AAB” for the embryo was defined as a maternal error (determined according to BAF around 0.33 and 0.66). The total count of paternal errors (Sp) and maternal errors (Sm) were used to compute a “Parental origin score”, defined as *log(Sm/Sp)*. Positive values of this score denoted the ploidy abnormality was due to a maternal error, while negative scores indicated the ploidy abnormality was due to a paternal error.

To determine the meiotic phase of origin, the variant had to be within 5cM from the centromere (to avoid dissociation of the SNP from the centromere due to meiotic recombination events) and one parent (the one in whom the error had happened) had to be heterozygous (e.g., AB), while the other was homozygous (e.g., AA or BB). This information can be used to identify SNPs indicative of the presence of both parental homologs (BPH) or of the presence of a single parental homolog (SPH). Due to inherent genetic constraints, meiotic phase analysis could only be performed on triploid samples. Similarly, to the “Parental origin score”, a “Meiotic origin score” was computed as log(S2/S1), with S1 and S2 representing the observed number of BPH SNPs and SPH SNPs, respectively. As an example, if the maternal genotype was “AB” and the paternal genotype was “AA” at the same locus, an occurrence of “AAB” for the embryo was defined as BPH and thus meiosis I error, whereas “AAA or ABB” for the embryo were defined as a SPH and a meiosis II error. Positive values of this score denoted an enrichment in SPH and an MII origin of the extra set of chromosomes, while negative scores indicated a MI origin of the extra chromosomes set with most SNPs being BPH.

### Recombination analysis

Data from 41 triploid embryos with concurrent aneuploidy and monogenic disorder testing were used, along with the genotyping data from their parents, to estimate the rate of recombination.

For each embryo, autosomal SNPs were selected using the filtering parameters as previously described.

For each SNP, concordance with a BPH or SPH model was evaluated using probabilities according to Mendel’s first law. If the embryonic genotype was concordant with either the presence of BPH or SPH, a concordance score of 0 was assigned. Conversely, if the SNP was not concordant with the model, a score of 1 was assigned. For each SNP a delta of the BPH and SPH concordance score was calculated (0, 1, −1). Using a sliding window approach from telomere to telomere for each chromosome, an average of the score was computed for each window of 3 consecutive SNPs. A switch of the average concordance score from 1 to –1 (or vice versa) was considered as a marker of recombination due to evidence of a phase change from BPH to SPH (or vice versa). Therefore, this locus, and the closest preceding SNP with an opposite average concordance score value, were used as ending and starting points of genomic windows in which recombination occurred (Supplementary Table 5). The size of the sliding windows was chosen to account for embryo genotypes that could be attributed to both BPH and SPH and based on an estimation of the SNPs genotyping error rate (0.42%) calculated as an average of inconsistent embryo genotypes according to the parental genotypes (Supplementary Table 6). Therefore, the chance of observing three consecutive SNPs genotyping errors was as low as 7e-8 (0.0042^3).

### Genome-wide uniparental isodisomy in diploid blastocysts

To determine the incidence of diploid embryos with duplication of one and concomitant loss of the other parental genome (isodiploidy) we used a dataset of 74,009 embryo biopsies with a 46,XX karyotype from aneuploidy testing by targeted NGS (DATABASE D). While some samples may overlap with the first dataset of 96,660 biopsies, this dataset also includes additional clinical data obtained after the completion of the initial set of analysis on ploidy abnormalities.

A bespoke algorithm was developed to interrogate the overall fraction of heterozygous SNPs (Total het fraction, including heterozygous SNPs mapping on autosomes and on chromosome X) and the pericentromeric fraction of heterozygous SNPs (pericentromeric het fraction-within 10Mb of centromeres) of autosomes and chromosome X. Heterozygous fraction was defined as the fraction of SNPs with a BAF between 0.1 and 0.9. Samples with a total het fraction lower than the average for all samples and additionally with a statistically significant lower pericentromeric het fraction, compared to the average and to the overall fraction of the sample itself, were marked as possible recombinant isodiploidy cases. Subsequently, each sample was manually checked to confirm the partial genome-wide loss of heterozygosity as visible in the alternate allele fraction distribution in the aneuploidy testing plot. Genetic similarity via fingerprinting analysis was also performed to evaluate the genetic correlation between sibling embryos and, where available, parental DNA.

### Statistical methods

Association between ploidy abnormalities and maternal and/or paternal age was assessed by performing multivariate logistic regression. As a positive control, the same dataset was also analyzed to corroborate an association between advancing maternal age (grouped according to SART classification) and aneuploidy rates utilizing a Chi-square Test. Additionally, an independent dataset of 93,341 oocytes from 15,851 cycles was used to assess age association with morphologically atypical pronuclei patterns, which are believed to correlate with abnormal ploidy status. Validation of the parental age correlation findings was performed by analyzing an independent dataset of 44,324 embryos utilizing the same statistical methodology.

To study the recurrence of abnormalities, female and male patients were first grouped into five categories according to age, as described above. For each age category, the recurrence of ploidy abnormalities in the same IVF cycle was investigated by taking as a null model, a random permutation of the embryo ploidy level across all samples under analysis, ensuring the null model had the same distribution of the number of embryos per cycle as in the original dataset. A P-value was estimated for each observed recurrence in each age category as a cumulative probability from the corresponding null model. P-values were considered statistically significant if P<0.05 following Bonferroni correction.

Finally, T-test was used to test for correlation between recombination rates and average aneuploidy rate per embryo.

### Validation of genotyping methods

Validation of the genetic findings was performed by analysis of rebiopsies of 9 selected embryos (Supplementary Table 7). Reproducibility of our methods was assessed by analyzing new biopsies with the same technology and PGT algorithms. Additionally, the meiotic origin of triploidy, the number of recombination events in triploid embryos, and the diagnosis of isodiploidy were confirmed by means of SNP-array (Infinium™ Global Screening Array-24 v3.0 – Illumina).

For the SNP-array data, heterozygous SNPs for one parent and homozygous for the other were considered informative (i.e., AB and AA or BB). Assuming from previous PGT results that all abnormalities were maternal in origin, only maternal heterozygous SNPs were evaluated. Each embryonic SNP was assigned, in a binary fashion, a value of either “1” corresponding to BPH/SPH (heterozygous SNP, which includes genotypes such as “AAB” or “ABB”) or “0” corresponding to SPH (homozygous SNP). Evaluation of pericentromeric SNPs was performed to infer meiotic phase of origin.

To identify breakpoints of meiotic recombination in triploid embryos, genomic regions of BPH and SPH were defined using the moving average smoothing algorithm. A sliding window of 50 SNPs was applied from the centromere towards each telomere of each chromosome and the average of BPH and SPH SNPs value was calculated within the window.

BPH was assigned if at least 90% of SNPs within the window were value 1. Similarly, SPH was assigned if the SNPs within the window were equally distributed between 1 and 0 (threshold 50%). The assignment of a recombination breakpoint was made at the point of switch between BPH and SPH defined by these thresholds.

To determine the statistical significance of obtaining the observed overlap between recombination breakpoints defined by SNP-array and PGT-derived windows of recombination, a permutation approach was adopted for each embryo. For each PGT-derived recombination window, we obtained a window of the same size, but placed randomly on the same chromosome. This procedure was repeated 1,000 times and the overlap of SNP-array points, with the randomly placed windows, was computed to obtain a null distribution. P-values were estimated as cumulative probabilities of the observed overlap from the null distribution.

Seven out of seven triploid embryos, and two out of two isodiploid embryos were confirmed utilizing the same technology, with a 100% concordance rate (95%CI: 66.4-100.0%).

Complete lack of recombination was confirmed in the re-biopsy of one embryo. Re-biopsy of one embryo with an average number of recombination events confirmed the presence of recombination, although with a slight difference (16 windows in the clinical biopsy vs 21 windows in the re-biopsy). Finally, the performance of our bespoke genotyping tools used in PGT was compared to SNP-array results obtained on re-biopsies of five embryos (Supplementary Figure 6). Triploidy and isodiploidy and their respective meiotic origins were confirmed in all embryos by means of SNP-array. Recombination events detected using SNP-array in triploid embryos mapped within the recombination windows detected using PGT in 37 out of 38 events (97.4%; 95% CI: 86.2-99.9%) with a probability of it being due to random chance of P<0.001 (permutation test).

## Supplementary

**Supplementary Table 1.** Description of the four datasets used in the study.

| **DATASET ID** | **# samples** | **Brief description** | **Objective** |
| --- | --- | --- | --- |
| DATASET A | 96,660 | Embryo biopsies analyzed by targeted NGS from 2PN derived blastocyst stage embryos from ICSI cycles from IVF clinics located in the US | Main dataset employed for the investigation of haploidy, triploidy, and related biological/clinical factors in human embryos |
| DATASET B | 44,324 | Embryo biopsies analyzed by targeted NGS from blastocyst stage embryos from ICSI cycles from IVF clinics located in Europe | Validation dataset employed to corroborate findings related to triploidy and haploidy from DATASET A |
| DATASET C | 93,341 | Zygotes obtained following ICSI with annotation of the number of pronuclei to assess fertilization | Validation dataset employed for the investigation of the relationship between maternal age and abnormal fertilization in human embryos |
| DATASET D | 74,009 | Female embryos biopsies analyzed by targeted NGS from blastocyst stage embryos from ICSI cycles from IVF clinics located in US | Selected female embryos from an expanded version of DATASET A employed for the investigation of recombinant isodiploidy in human embryos |

**Supplementary Figure 1.**
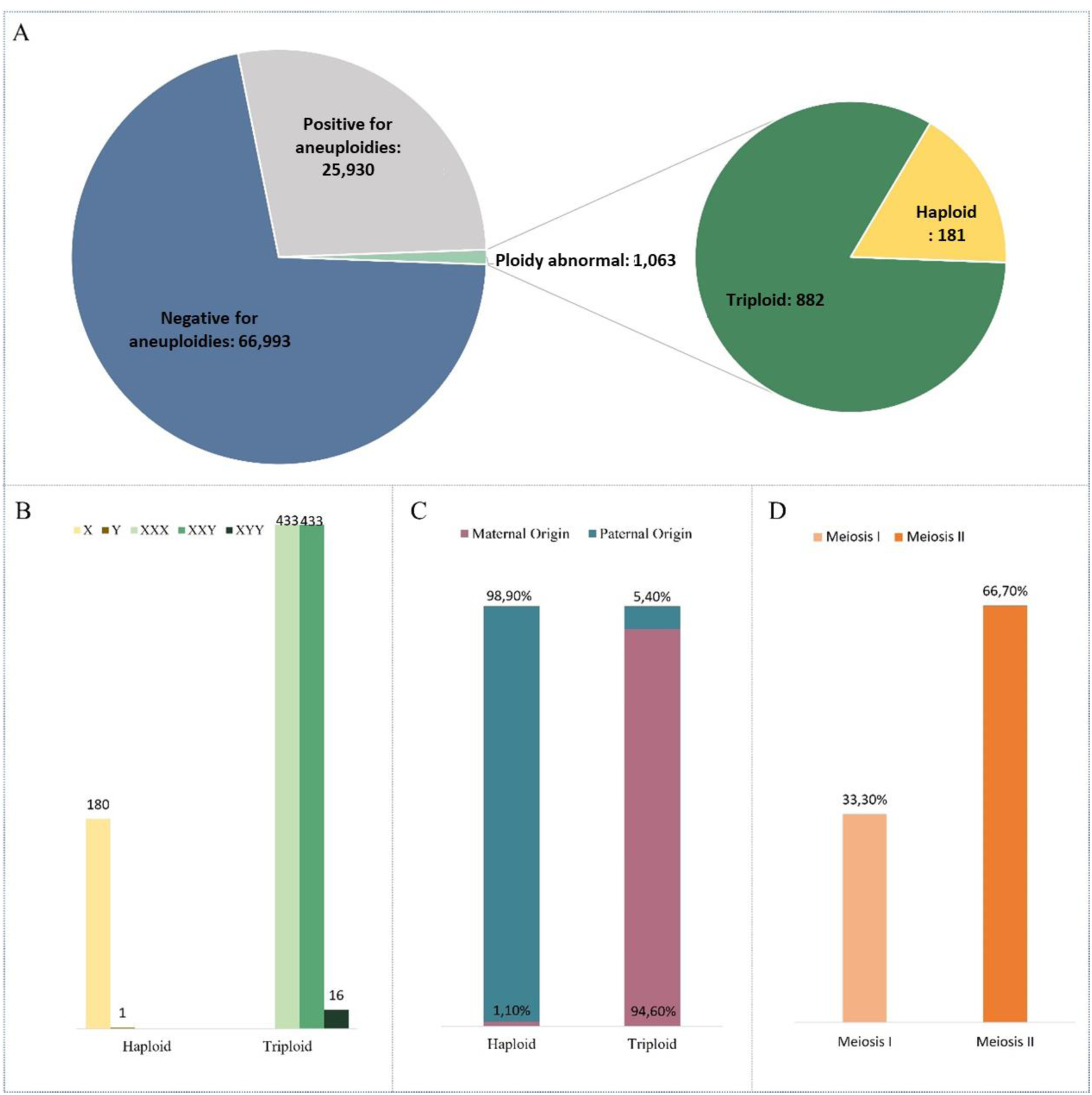
Overview of the diagnostic outcomes from preimplantation genetic testing. A: Pie chart illustrating the frequencies of aneuploidy, euploidy, haploidy, and triploidy diagnoses in preimplantation embryos. B: Histogram depicting the distribution of different sex chromosome combinations observed in haploid and triploid embryos. C: Histogram showing the percentage of haploid embryos, categorized by whether the chromosomal abnormality originated maternally or paternally. D: Histogram showing the meiotic origin of triploid embryos, distinguishing between segregation errors occurring in meiosis I versus meiosis II (without differentiating between male and female meiosis).

**Supplementary Figure 2.**
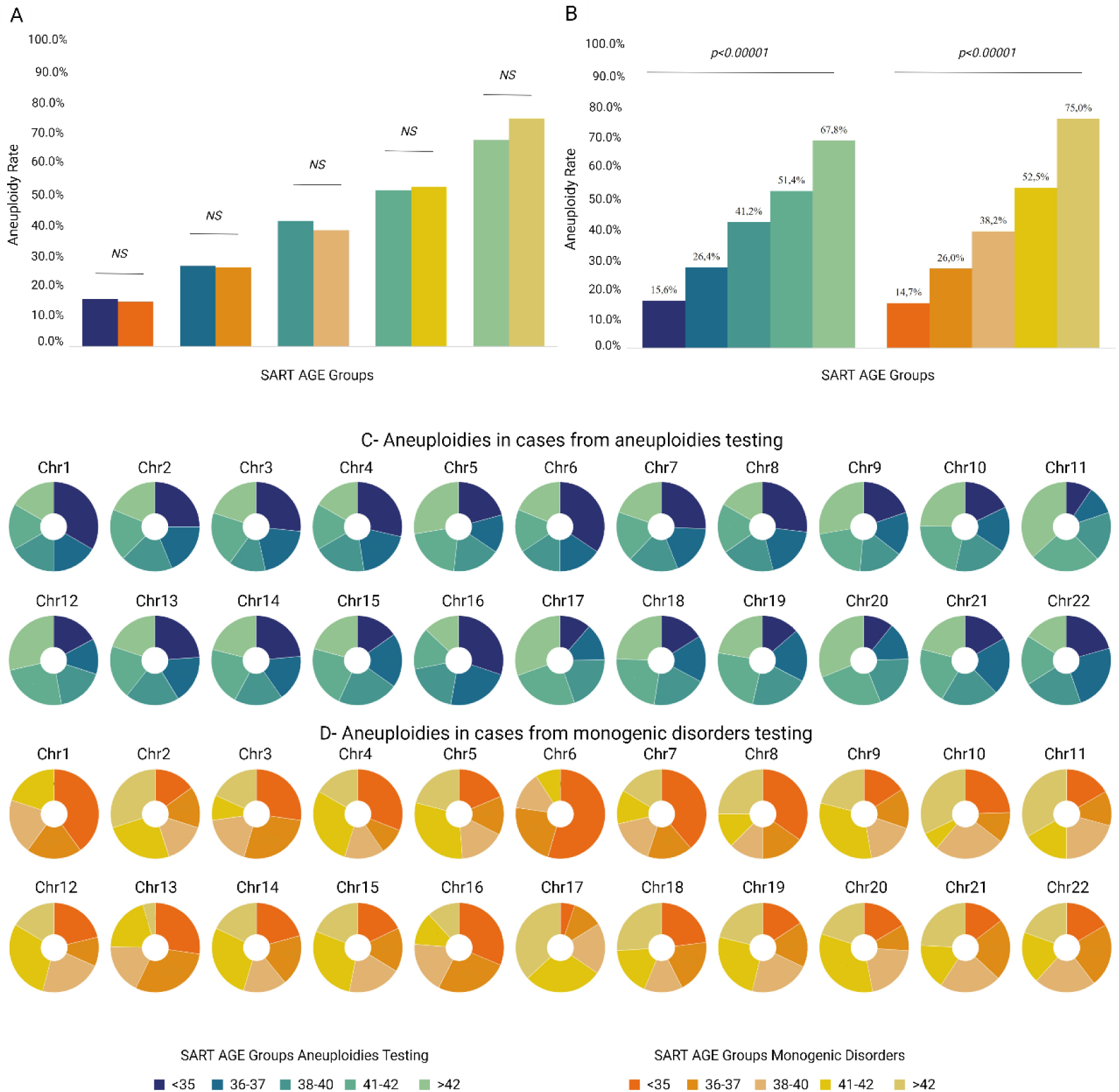
Impact of maternal age on aneuploidy rates across preimplantation genetic testing indications for datasets A and. **B.** A and B-Histograms depicting aneuploidy rates in the 5 SART age groups. No statistical differences were detected amongst the different SART age groups when comparing indication for testing. However, both groups demonstrated a statistically significant correlation between advanced maternal age and increased aneuploidy rates. C-D Pie charts showing the frequencies of aneuploidies (both trisomies and monosomies) for each autosomal chromosome according to SART age group.

**Supplementary Table 2.** Number of usable SNPs and detectable recombination events per each triploid embryo in the research pipeline.

| PGT-M_SAMPLE | NUMBER OF USABLE IN-FORMATIVE SNPs | RECOMBINATION EVENTS |
| --- | --- | --- |
| 1 | 1771 | 0 |
| 2 | 1745 | 0 |
| 3 | 1590 | 0 |
| 4 | 1583 | 0 |
| 5 | 1685 | 0 |
| 6 | 1473 | 0 |
| 7 | 1408 | 0 |
| 8 | 1737 | 8 |
| 9 | 1493 | 9 |
| 10 | 1668 | 10 |
| 11 | 1740 | 11 |
| 12 | 1708 | 13 |
| 13 | 1643 | 13 |
| 14 | 1701 | 13 |
| 15 | 1512 | 13 |
| 16 | 1623 | 13 |
| 17 | 1598 | 14 |
| 18 | 1513 | 14 |
| 19 | 1730 | 15 |
| 20 | 1785 | 16 |
| 21 | 1662 | 16 |
| 22 | 1723 | 16 |
| 23 | 1670 | 16 |
| 24 | 1745 | 17 |
| 25 | 1855 | 17 |
| 26 | 1690 | 17 |
| 27 | 1765 | 18 |
| 28 | 1765 | 18 |
| 29 | 1795 | 18 |
| 30 | 1666 | 18 |
| 31 | 1699 | 18 |
| 32 | 1615 | 19 |
| 33 | 1605 | 19 |
| 34 | 1708 | 20 |
| 35 | 1569 | 20 |
| 36 | 1355 | 20 |
| 37 | 1572 | 20 |
| 38 | 1584 | 21 |
| 39 | 1724 | 21 |
| 40 | 1642 | 21 |
| 41 | 1854 | 26 |

**Supplementary Figure 3.**
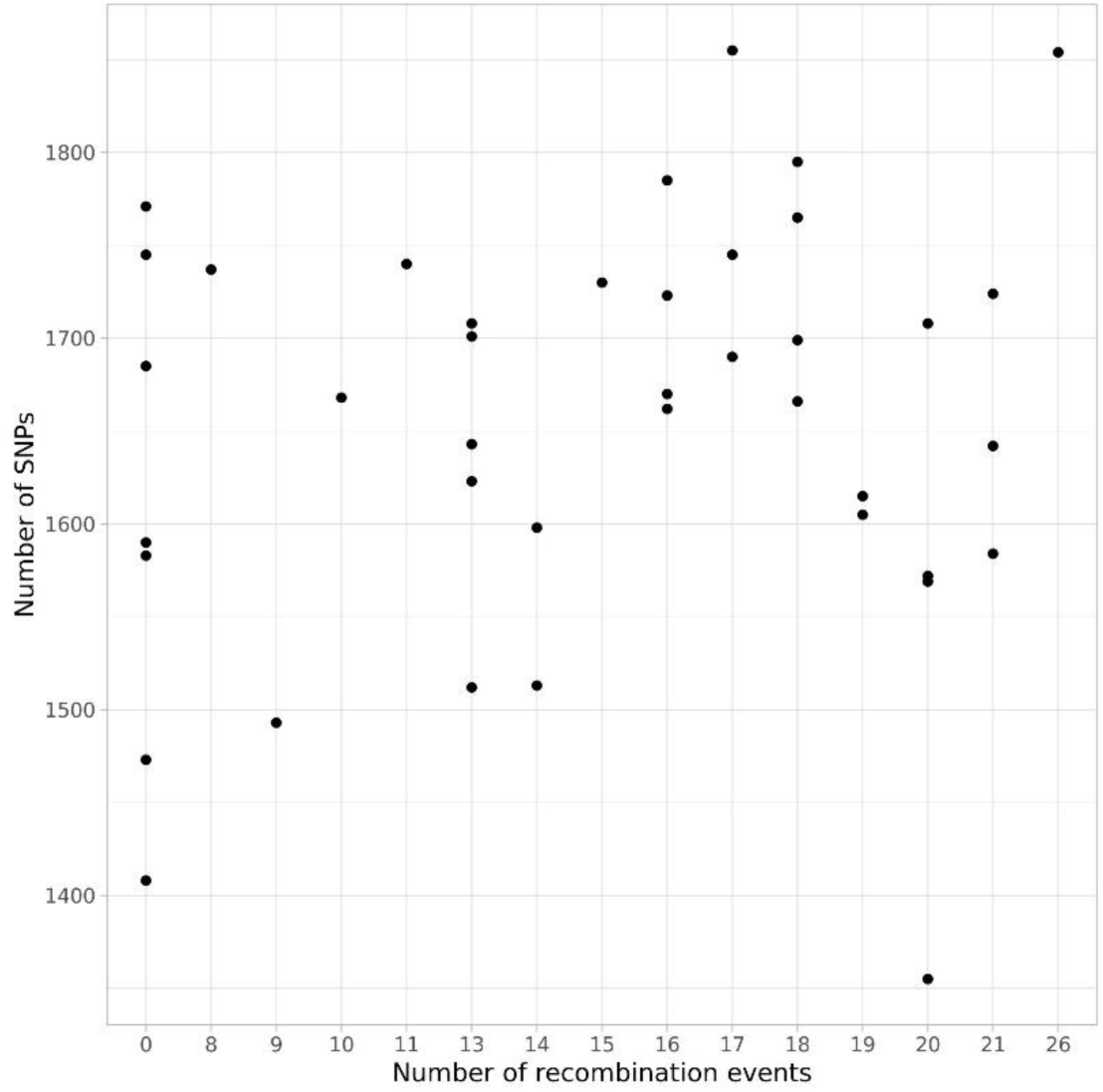
Distribution of the number of SNPs (y axis) vs the number of recombination events (x axis).

**Supplementary Table 3.** Paternal age is not correlated to embryonic ploidy, even when corrected for maternal age.

| Age groups | B | S.E. | p value | OR | 95% CI per OR |  |
| --- | --- | --- | --- | --- | --- | --- |
| paternal.age <35 yo |  |  | 0,587 |  |  |  |
| paternal.age 35-37 yo | -0,176 | 0,114 | 0,122 | 0,838 | 0,67 | 1,048 |
| paternal.age 38-40 yo | -0,156 | 0,125 | 0,212 | 0,856 | 0,67 | 1,093 |
| paternal.age.group 41-42 yo | -0,134 | 0,155 | 0,385 | 0,874 | 0,646 | 1,184 |
| paternal.age.group > 42 yo | -0,17 | 0,138 | 0,218 | 0,844 | 0,644 | 1,106 |
| maternal.age <35 yo |  |  |  |  |  |  |
| maternal.age.group 35-37 yo | 0,414 | 0,119 | **0,001** | 1,512 | 1,197 | 1,91 |
| maternal.age.group 38-40 yo | 0,503 | 0,118 | **0,000** | 1,654 | 1,311 | 2,085 |
| maternal.age.group 41-42 yo | 0,873 | 0,14 | **0,000** | 2,394 | 1,818 | 3,152 |
| maternal.age.group > 42 yo | 0,804 | 0,151 | **0,000** | 2,234 | 1,663 | 3,001 |

**Supplementary Figure 4.**
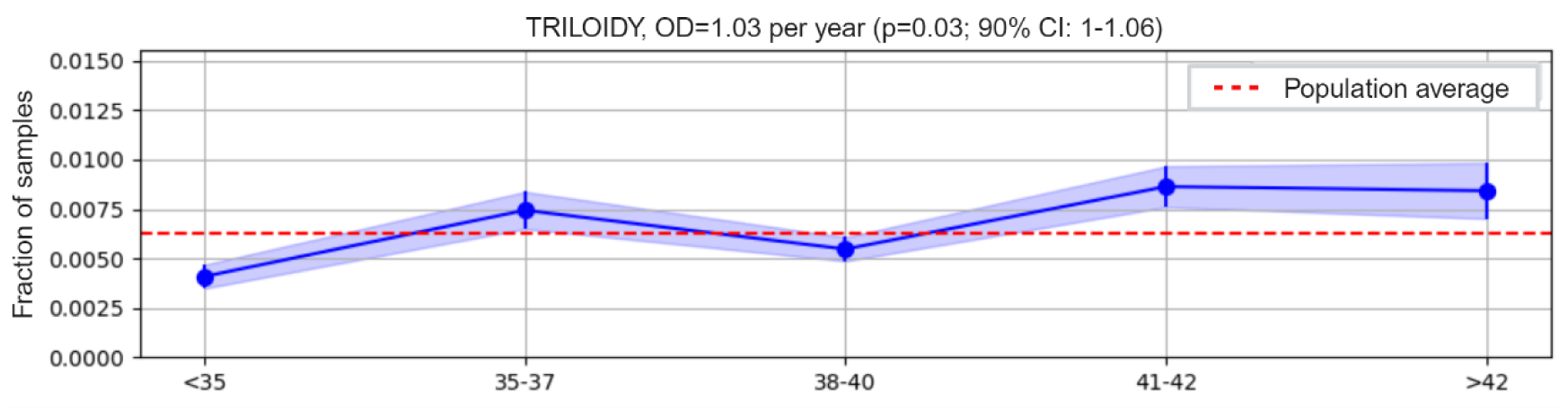
The risk of ploidy abnormalities rises with advancing maternal age: confirmation from an independent dataset. Plot illustrating the relationship between maternal age and triploidy in an independent dataset. Maternal ages were classified using the SART age groups (X axis). The Y axis shows the fraction of affected triploid embryos.

**Supplementary Figure 5.**
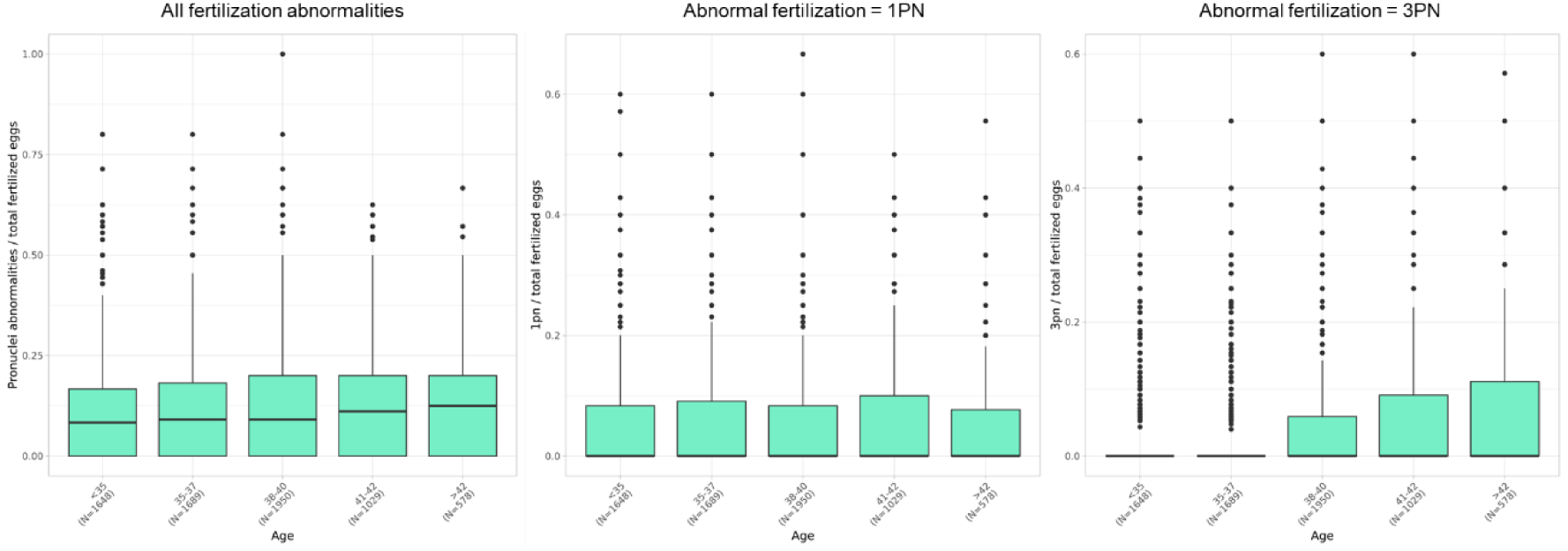
Maternal age is correlated with an increased risk of abnormal fertilization. Box plots representing, from left to right: distribution of (i) all abnormally fertilized oocytes, (ii) of oocytes with 1 pronucleus and (iii) of oocytes with 3 pronuclei, against female age groups. Only cycles with more than 5 inseminated oocytes are plotted.

**Supplementary Table 4.** Ploidy abnormalities recurrence (≥3). Six couples were found to have a ploidy abnormality recurrence within a single IVF/ICSI cycle.

| CoupleID_Embryo | Female Age Grouped | Male Age Grouped | Ploidy level |
| --- | --- | --- | --- |
| 1_1 | <35 | 35-37 | diploid |
| 1_2 | <35 | 35-37 | diploid |
| 1_3 | <35 | 35-37 | **triploid** |
| 1_4 | <35 | 35-37 | **haploid** |
| 1_5 | <35 | 35-37 | diploid |
| 1_6 | <35 | 35-37 | diploid |
| 1_7 | <35 | 35-37 | diploid |
| 1_8 | <35 | 35-37 | diploid |
| 1_9 | <35 | 35-37 | **triploid** |
| 2_1 | <35 | <35 | diploid |
| 2_2 | <35 | <35 | **triploid** |
| 2_3 | <35 | <35 | **triploid** |
| 2_4 | <35 | <35 | **triploid** |
| 3_1 | 41-42 | 38-40 | **triploid** |
| 3_2 | 41-42 | 38-40 | diploid |
| 3_3 | 41-42 | 38-40 | diploid |
| 3_4 | 41-42 | 38-40 | diploid |
| 3_5 | 41-42 | 38-40 | **triploid** |
| 3_6 | 41-42 | 38-40 | diploid |
| 3_7 | 41-42 | 38-40 | **triploid** |
| 4_1 | <35 | <35 | diploid |
| 4_2 | <35 | <35 | diploid |
| 4_3 | <35 | <35 | diploid |
| 4_4 | <35 | <35 | **triploid** |
| 4_5 | <35 | <35 | **triploid** |
| 4_6 | <35 | <35 | diploid |
| 4_7 | <35 | <35 | **triploid** |
| 4_8 | <35 | <35 | diploid |
| 5_1 | <35 | <35 | **triploid** |
| 5_2 | <35 | <35 | **triploid** |
| 5_3 | <35 | <35 | **triploid** |
| 6_1 | 35-37 | 35-37 | diploid |
| 6_2 | 35-37 | 35-37 | **triploid** |
| 6_3 | 35-37 | 35-37 | **triploid** |
| 6_4 | 35-37 | 35-37 | diploid |
| 6_5 | 35-37 | 35-37 | diploid |
| 6_6 | 35-37 | 35-37 | diploid |
| 6_7 | 35-37 | 35-37 | **triploid** |

**Supplementary Table 5.** Examples explicative of the scoring system used for probabilities and inconsistencies of BPH and SPH. Given specific allelic combination in the parents, for each embryonic genotype a probability of it BPH or SPH was assigned. Subsequently, each probability equal to 0 and 1 was converted into an inconsistency score of 1 or 0, respectively.

| Female<br>genotype | Male<br>genotype | Embryo<br>genotype | Probability<br>BPH | Probability<br>SPH | Inconsistency<br>BPH | Inconsistency<br>SPH |
| --- | --- | --- | --- | --- | --- | --- |
| AB | AA | AAA | 0 | 1 | 1 | 0 |
| AB | AA | ABB | 0 | 1 | 1 | 0 |
| AB | AA | AAB | 1 | 0 | 0 | 1 |
| AB | BB | BBB | 0 | 1 | 1 | 0 |
| AB | BB | AAB | 0 | 1 | 1 | 0 |
| AB | BB | ABB | 1 | 0 | 0 | 1 |

**Supplementary Table 6.** Evaluation of genotyping error rate in triploid embryos. Given specific allelic combination in the parents, embryonic genotype at that locus had to be either heterozygous or homozygous. If embryonic genotype was inconsistent due to allele drop out or allele drop in mechanisms, this was interpreted as indication of a SNP genotyping error.

| Female genotype | Male genotype | Obligated embryonic genotype | Inconsistent embryonic genotypes | Meaning |
| --- | --- | --- | --- | --- |
| AA | BB | AAB or ABB (hetero) | AAA and BBB (homo) | Allele Drop Out (ADO) |
| BB | AA | BBA or BAA (hetero) | AAA and BBB (homo) | Allele Drop Out (ADO) |
| AA | AA | AAA (homo) | AAB or ABB (hetero) | Allele Drop In (ADI) |
| BB | BB | BBB (homo) | AAB or ABB (hetero) | Allele Drop In (ADI) |

**Supplementary Table 7.** Schematic summary of biological and analytical validation of the genotyping tools.s.

|  | Biological validation (1st Rebiopsy) |  |  | Analytical validation (2nd Rebiopsy) |  |
| --- | --- | --- | --- | --- | --- |
| Original PGT Results | Re-biopsy for confirmation of ploidy abnormality using PGT | Re-biopsy for meiotic origin and recombination using PGT | Origin | Re-biopsy using SNP-array | Origin |
| Triploid | yes | yes | TE | yes | TE |
| Triploid | yes | yes | TE | yes | TE |
| Triploid | yes | NA | ICM | yes | TE |
| Triploid | yes | NA | ICM | NA | - |
| Triploid | yes | NA | ICM | NA | - |
| Triploid | yes | NA | ICM | NA | - |
| Triploid | yes | NA | ICM | NA | - |
| Isodiploid | yes | NA | TE | yes | TE |
| Isodiploid | yes | NA | TE | yes | TE |

**Supplementary Figure 6.**
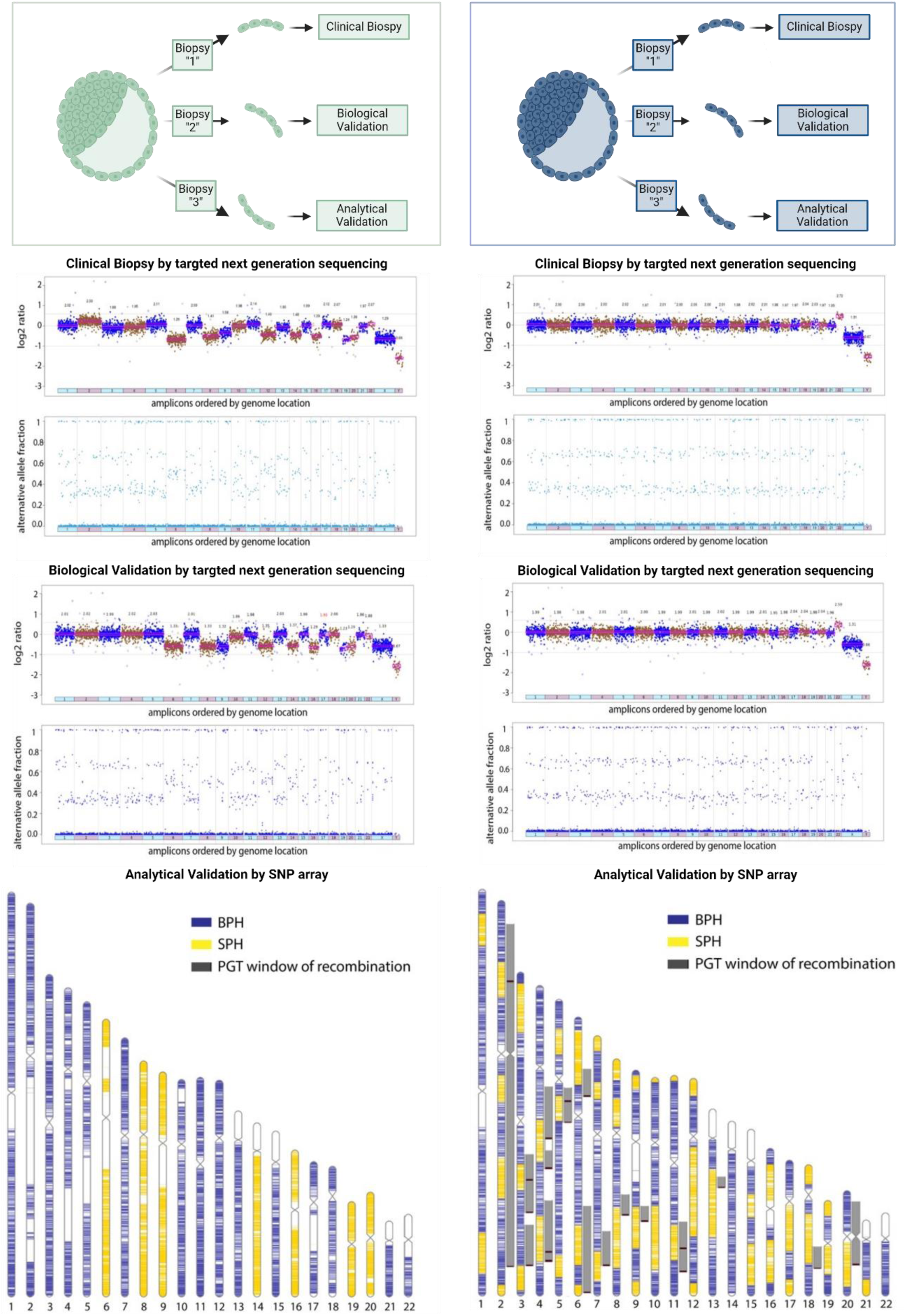
Reproducibility of the methods and concordance with SNP-array. This figure illustrates the multi-biopsy model, where three consecutive trophectoderm biopsies were taken from a single embryo. The first biopsy served as a clinical sample for preimplantation genetic testing (PGT), the second for biological validation, and the third for analytical validation using SNP-array analysis. The left panel depicts an embryo with lack of recombination, meiosis I origin and additional aneuploidies. The panel on the right depicts an embryo with average recombination rate, meiosis I origin of triploidy and one additional aneuploidy on chromosome 22. The first two plots (top to bottom) show the concordance between the clinical biopsy (1^st^ Biopsy) and the rebiopsy (2^nd^ Biopsy) in terms of ploidy level abnormalities and additional aneuploidies. The last plot shows ideograms of the maternally inherited chromosomes distribution of SPH (yellow) and BPH (blue) SNPs according to SNP-array. The white areas on the chromosome ideograms indicate regions with no SNP coverage. Grey bars adjacent to the ideograms (separated by small black bars where two grey bars meet at the same genomic region) depict PGT windows of recombination, within which the switches from BPH to SPH were identified.

## Supplementary Note 1

## A method to infer the total map length considering that the proportion of chiasmata that are visible as a crossover is different between the MI-error-like state (both parental homologs) and MII-error-like state (same parental homolog)

As shown in the illustration at the end of this text, any chiasma in a single parental homolog (SPH) segment will change the state to both parental homolog (BPH). In contrast, only half of the chiasmata in BPH segments will be observed as changing the state to SPH. We assume no crossover interference and no chromatid interference. (Such that each chiasma has an independent probability of ½ to be observed.) Denote the chiasmata rate along the genome (say, per bp) as *λ*. We assume the same rate regardless of whether the state is SPH or BPH.

For an SPH segment of length *x*, the probability of no event up to length *x* and then transition to BPH is *λe*^−*λx*^.

For a BPH segment of length *x*, the probability of no event up to length *x* and then transition to SPH is 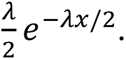 This is because the probability of a transition (in each infinitesimal length unit) is *λ* for a chiasma, and then ½ for the chiasma to be observed.

Suppose we have *L*_*s*_bp in SPH state and *L*_*b*_in BPH, with *L* = *L*_*s*_ + *L*_*b*_. Denote the number of events at the end of SPH and BPH segments as *n*_*s*_ and *n*_*b*_, respectively, and *n* = *n*_*s*_ + *n*_*b*_.

The likelihood of the data is thus: 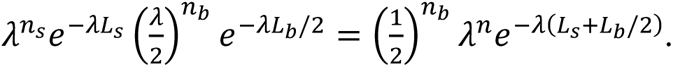

The log likelihood is (up to factors independent of *λ*) *n* log *λ* − *λ*(*L*_*s*_ + *L*_*b*_/2). Taking the derivative with respect to *λ* and equating to zero gives an equation for the maximum likelihood estimator,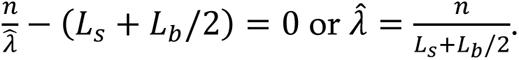

This gives an estimate for the rate in which chiasmata are generated along the genome. The estimated total number of chiasmata genome-wide is 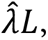, and the estimated total map length, denoted 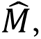 is 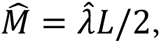 because only half the chiasmata become crossovers. The expected number of crossovers genome-wide is the total map length.

We thus have 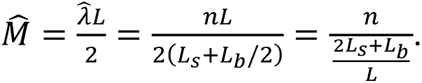

Finally, denote by *α* = *L*_*s*_/*L* the proportion of the genome in SPH segments. Accordingly, *L*_*b*_/*L* = 1 − *α*. We thus have 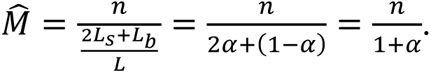

This makes sense, because, for example, if *α* ≪ 1 (almost all BPH), we have 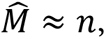 because we observe only half the chiasmata to begin with, which is the expected number of crossovers. If *α* ≈ 1 (almost all SPH), we have 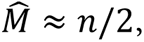 because we observed every single chiasmata, so we must divide by 2 to get the number of crossovers.

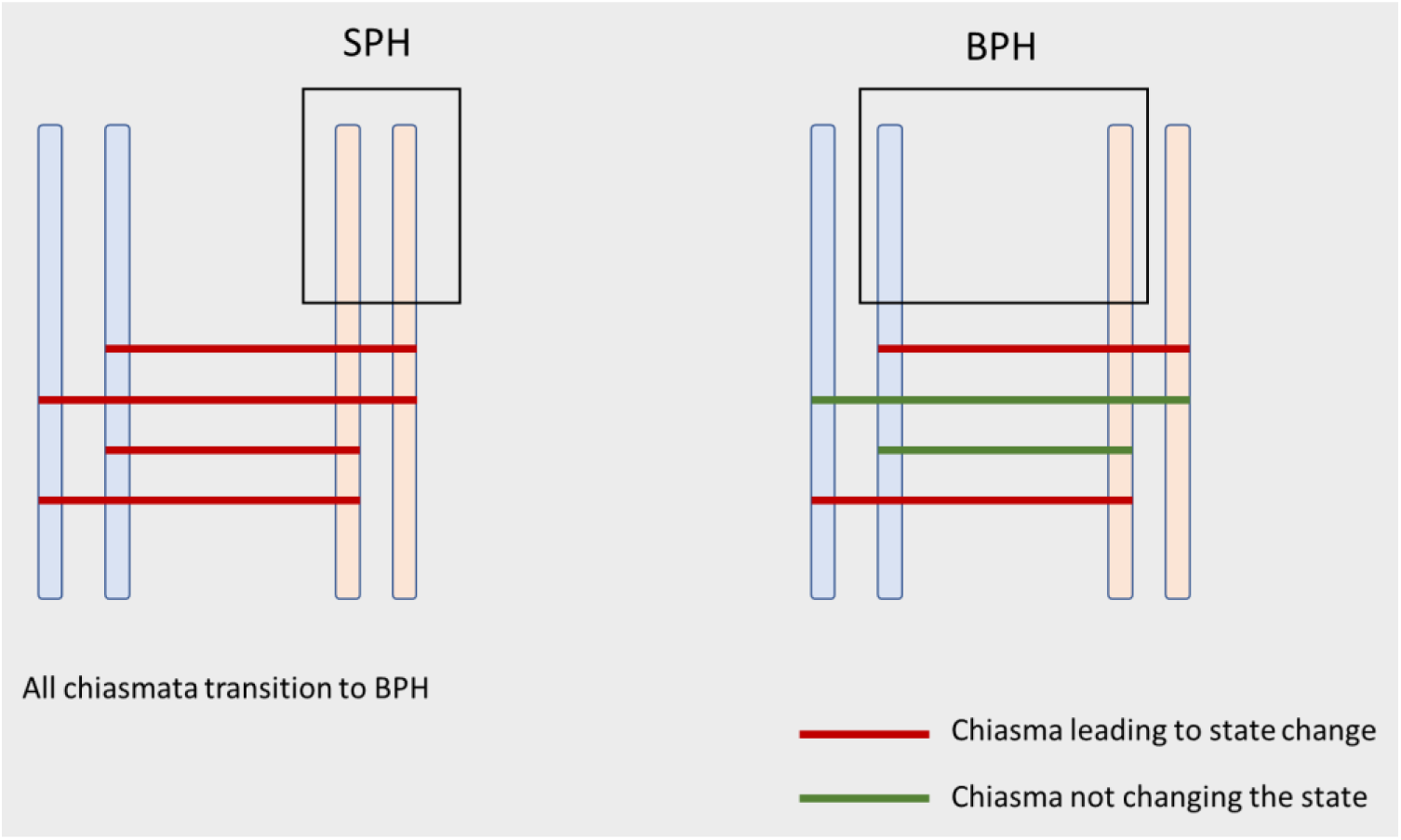

